# The intergenerational transmission of educational attainment: A closer look at the (interrelated) roles of paternal involvement and genetic inheritance

**DOI:** 10.1101/2022.04.06.487369

**Authors:** Renske Verweij, Renske Keizer

## Abstract

Numerous studies have documented a strong intergenerational transmission of educational attainment. In explaining this transmission, separate fields of research have studied separate mechanisms. To obtain a more complete understanding, the current study integrates insights from the fields of behavioural sciences and genetics and examines the extent to which factors from each field are unique underlying mechanisms, correlate with each other, and/or act as important confounders in the intergenerational transmission of educational attainment. Specifically, we focus on paternal involvement as our behavioural indicator and children’s polygenic score (PGS) for education as our genetic indicator and assess the role that fathers play in the intergenerational transmission of educational attainment. To answer our research questions, we use rich data from The National Longitudinal Study of Adolescent to Adult Health (n=4,579). Firstly, results from our mediation analyses showed that about 4 per cent of the effect of fathers’ educational attainment on children’s educational attainment is explained by paternal involvement, whereas a much larger share, 21 per cent, is explained by children’s education PGS. Secondly, our results showed that these genetic and behavioural influences are significantly correlated to each other. Thirdly, we found support for genetic confounding, as adding children’s education PGS to the model reduced the effect of paternal involvement by 11 per cent. Fourthly, evidence for social confounding was almost negligible (a reduction of half a per cent of the genetic effect). Our findings highlight the importance of integrating insights and data from multiple disciplines in understanding the mechanisms underlying the intergenerational transmission of inequality, as our study reveals that behavioural and genetic influences overlap, correlate, and confound each other as mechanisms underlying this transmission.

## Introduction

Individuals with highly educated parents generally achieve better outcomes in school than individuals with lower educated parents (1). For example, during primary and secondary school years, children with highly educated parents obtain higher school grades (2), and eventually, children with at least one highly educated parent are more than twice as likely to obtain tertiary education themselves compared to children without highly educated parents (3). Different research fields focus on different factors in explaining this intergenerational transmission of educational attainment. Research from the field of behavioural sciences aims at explaining this intergenerational transmission by considering, amongst others, characteristics of the family environment, such as parenting practices and the level of social, cultural, and economic capital within families (4, 5). In contrast, research from the field of genetics focuses on the role of genetic transmissions (6).

With a few exceptions (7–10), studies that have looked at the *interrelated* linkages between behavioural and genetic factors are rare. That such research is rare is very unfortunate; investigating behavioural and genetic factors in isolation most likely overestimates the unique contribution each factor makes. In line with this idea, previous studies have revealed that controlling for genetics reduced the impact of parenting quality (8) and family environment (10) on children’s educational achievement, which indicates that part of the assumed social effect might be genetic, so-called ‘genetic confounding’. Research has also shown the reverse; part of the genetic effect on education might be socially confounded; empirical studies have revealed that the influence of parents’ and children’s education-related genes on children’s educational attainment is reduced when parenting practices and family SES are taken into account (9–11).

These findings underscore the importance of taking both behavioural and genetic factors into account to obtain a clear understanding of the mechanisms underlying the intergenerational transmission of educational attainment, which is the aim of the current paper. Our study builds forward on work by Wertz et al (2020). Although these authors did not investigate the intergenerational transmission of educational attainment, they did scrutinize the role of maternal behaviour and genes in explaining children’s educational attainment. Their study revealed that mothers’ cognitively stimulating parenting explained the effect of mothers’ education polygenic score (PGS) on children’s educational attainment and that the inclusion of children’s education PGS slightly reduced the effect of mothers’ parenting (cognitive stimulation, warmth, and sensitivity) on children’s education attainment (8). In the current paper, we turn our attention to the role of fathers (whilst controlling for the role mothers play).

Traditionally, studies on the intergenerational transmission of education/socioeconomic status (SES) focused on obtaining information on fathers’ educational attainment or profession. As women, particularly in previous decades, often retreated from the labour market after marriage or childbirth, information on mother’s educational attainment or profession was not always available for each child. In contrast, information on the role fathers played in parenting was often neglected or overlooked, as mothers were the primary source of information to report on child development. During the last 50 years, however, fathers have become more and more involved in parenting (12, 13). Although some scholars argue and show that certain roles might still be most prominent amongst solely mothers or solely fathers, the roles of fathers (and mothers alike) are increasingly being expanded (14), which have made fathers and mothers more similar in their roles as caregivers (15). Even though some scholars construct a gender-differentiated vision of *how* mothers and fathers parent, there is increasing consensus among scholars that there are little to no differences in how *well* mothers and fathers parent (e.g. Fagan *et al.* 2014). In line with this, and pertaining to educational attainment, studies show that the relationship between parental involvement and academic achievement is similar for mothers and fathers (16).

That said, we do expect to see greater variation in paternal than in maternal involvement, and this is our main rationale for choosing to focus on paternal involvement in the current paper. Over the years, fathers in two-parent families have spent more time with their children (17), a pattern especially common among higher educated fathers (18). After divorce, which is more common amongst lower-educated families (19), fathers often are, or become, less involved in their children’s lives, which is especially the case among lower-educated families (20). This greater variation in father involvement across social strata suggests that father involvement could be an important underlying mechanism of the intergenerational transmission of educational attainment, and thus possible leverage to reduce inequality.

Several studies have investigated the role that paternal involvement plays in children’s educational attainment and in the intergenerational transmission of educational attainment (e.g. Gordon, 2017; Whitney *et al.* 2018). Unfortunately, however, these studies did not control for genetic effects. Even though twin studies show that both genetic aspects, as well as the shared environment, are important in explaining children’s educational attainment, the shared environment in these studies remains unmeasured (e.g. Branigan, McCallum, & Freese, 2013; Johnson, Deary, & Iacono, 2009; Nielsen, 2006; Schulz, Schunck, Diewald, & Johnson, 2017).

In sum, the current study aims to provide a clearer understanding of the (interrelated) roles that paternal involvement and genes play in the intergenerational transmission of educational attainment. We use children’s PGS for educational attainment as our genetic indicator. This PGS is based on a genome-wide association study (GWAS) conducted among 1.1 million individuals (26). These GWAS summary statistics are used to calculate the sum of all risk alleles, weighted by their reported effect sizes. A PGS thus can be seen as the summary measure of the genetic propensity for a trait (27). The Education PGS has been found to explain about 11-13% of the variation in educational attainment (26).

## Theory and hypotheses

### Paternal involvement as an independent mechanism underlying the intergenerational transmission of educational attainment

To build the argument that fathers’ involvement in their children’s lives is an underlying mechanism for the intergenerational transmission of educational attainment, we would first have to argue and show that (1) fathers’ educational attainment is significantly associated with children’s educational attainment, that (2) fathers’ involvement is significantly associated with children’s educational attainment and (3) fathers’ educational attainment is significantly associated with fathers’ involvement in their children’s lives (see Figure 1).

**Fig 1.**
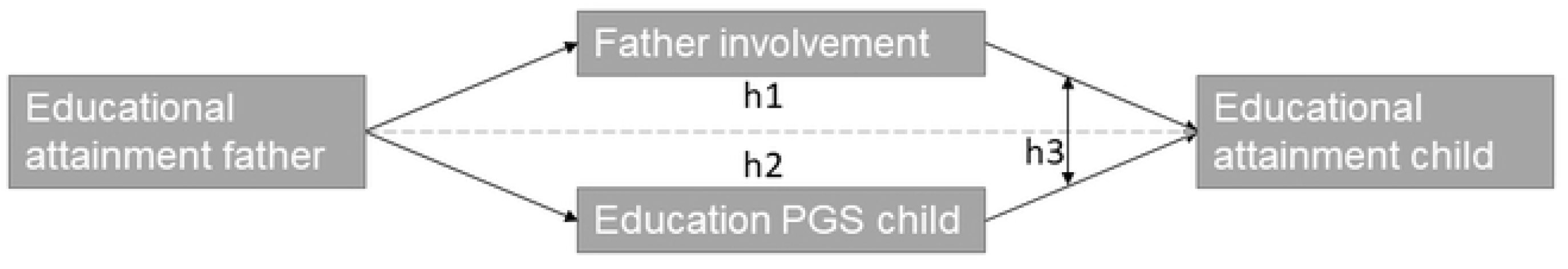
Graphical presentation of hypotheses 1, 2, and 3: mediation pathways and correlated effects.

Firstly, numerous studies have revealed that fathers’ educational attainment influences children’s cognitive functioning (see for a review Varghese & Wachen 2016). Children of highly educated fathers not only show better cognitive functioning but also obtain better school grades throughout their educational career (2) and higher educational levels (3). Secondly, fathers’ involvement in their children’s lives has been argued to positively affect their children’s educational attainment. Different dimensions of fathers’ involvement might uniquely influence children’s educational performances. First of all, and most directly, fathers can influence their children’s educational outcomes through their involvement in teaching-related activities such as coaching, helping with homework, communication with school personnel, and active participation in classroom or school activities (21,29,30). The involvement of fathers may benefit children’s school achievement, amongst others because the help fathers provide and the time that fathers practice with their children can improve their children’s skills (31). In addition, the greater involvement of fathers in schoolwork could positively impact children’s educational achievements, because greater school involvement of parents results in more ambitious school aspirations, greater motivation to perform well in school, and higher school attendance (32). The rationale behind these findings is that greater involvement of parents is indicative of the value they attach to educational achievements, and children internalize these values (31). Previous research found that children who have fathers that are more involved in their schoolwork obtained higher grades in high school (21,29,30). These positive associations remain when controlling for the involvement of the mothers (21, 30). Besides fathers’ involvement in school (work), the frequency of joint activities undertaken by father and child is also shown to benefit children’s educational attainment (33, 34). Previous studies show that children who spend more time alone with their father score higher on cognitive tests (33) (also when controlling for the involvement of the mother) and that the more time fathers spend with their children at young ages on care-related tasks as well as playing and reading, the better children fare in school tasks (34). Thirdly, there is empirical evidence showing that fathers’ educational attainment is related to fathers’ involvement (18). Highly educated fathers are generally more involved in their children’s lives, amongst others because these fathers feel more confident in helping their children with schoolwork (35, 36), because they have more intensive parenting ideologies (5) and they have more time available (18).

Based on the theoretical and empirical work, it is likely that fathers’ involvement mediates the association between fathers’ and children’s educational attainment. Our first hypothesis therefore reads: *Fathers’ involvement (school-specific involvement and leisure involvement) is an underlying mechanism for the impact of fathers’ educational attainment on children’s educational attainment (H1).* A small number of studies have shown that fathers’ involvement indeed serves as a mediator between educational attainment and child outcomes (29,37–39) A clear limitation is that they did not control for children’s genetic characteristics, and therefore might not have been able to obtain a reliable understanding of the role father involvement plays in the intergenerational transmission of educational attainment.

### Genetic influences as an independent mechanism underlying the intergenerational transmission of educational attainment

A large number of twin studies showed that educational attainment is approximately 40% heritable, that is, 40% of the variation in education can be explained by genetic variation (see the meta-analysis by Branigan et al., 2013). This heritability can be explained partly by the heritability of intelligence, but also by the heritable component of amongst others personality traits, self-efficacy, and behavioural problems (40). Children with genes that are positively related to higher educational attainment are more open, agreeable, conscientious, and show more academic motivation, which all result in better educational achievements (41). As parents and children are 50% genetically similar, part of the intergenerational transmission of education is posed to be through genetic influences. Several previous studies showed that genes can explain part of the intergenerational transmission of education (42–44). Based on the above-described literature we expect to find that *genetic influences are an underlying mechanism for the impact of fathers’ educational attainment on children’s educational attainment (H2)*.

### Three types of correlations between genetic influences and father involvement

There are three different reasons why we can expect to find a correlation between father involvement and children’s education PGS. These three reasons are related to the three types of rGE that are theoretically distinguished (45). The first type is an *evocative gene-environment correlation*: parents will behave differently to a child based on the child’s characteristics (which are genetically influenced) (46). In our case, children’s education PGS does not only capture the intelligence of the child but also personality traits, academic motivation, behavioural problems, and self-efficacy (40, 41). It is therefore likely that children with a higher education PGS more often evoke their father’s involvement (in particular with school), as these children are also more likely to be more interested in learning. If this is true, this would result in a positive rGE.

The second type is an *active gene-environment correlation*: children with higher education PGS might not only be more likely to be more interested in learning, but they might also be more prone to actively seek help from their parents with homework or discuss school matters with their father, which then results in greater involvement of the father.

The third type is a *passive gene-environment correlation*: children inherit half of their genes from each parent, and these parents also rear them and shape their environment. This can result in a correlation between the child’s education PGS and father’s involvement in two ways. Firstly, children with a high education PGS are more likely to have a parent with a high education PGS, and the parent’s education PGS not only shapes their own educational attainment but also their parenting practices (9). Secondly, children with a higher education PGS generally have a highly educated father, and because of reasons such as status maintenance motives (47), these highly educated fathers are more likely to be more involved in their children’s lives.

Based on the abovementioned theoretical considerations, *we expect that father involvement (school-specific involvement and leisure involvement) is positively correlated to children’s education PGS in their relation to education (H3).* Several previous studies showed a correlation between genes and different aspects of the family environment, such as parental sensitivity, warmth, stimulating parenting, and parental SES (7–9,48), and one study looked specifically at the correlated effect of genes and family environment on education (10). No study has looked at the correlation between genes and father involvement in their relation to education.

### Implications of correlations between genetic factors and father involvement for understanding the mechanisms underlying the intergenerational transmission of educational attainment: Genetic confounding

In the context of the expected correlations between genetic factors and father involvement, it might be the case that part of the effect of father involvement is driven by genetic factors, so-called genetic confounding; both fathers’ involvement as well children’s educational attainment is shaped by the same genetic factors (see Figure 2 for a graphical representation). This is due to the passive rGE. For example, the same genes that result in higher education, also result in greater involvement of fathers through personality traits or a sense of responsibility of the father. Previous research by Wertz and colleagues showed that controlling for genetics reduced the effect of parental warmth and sensitivity on the child’s educational achievement to some extent (8). In line with this finding, we expect that *the genetic influences partly explain the behavioural mechanism underlying the intergenerational transmission of educational attainment (H4)*.

**Fig 2.**
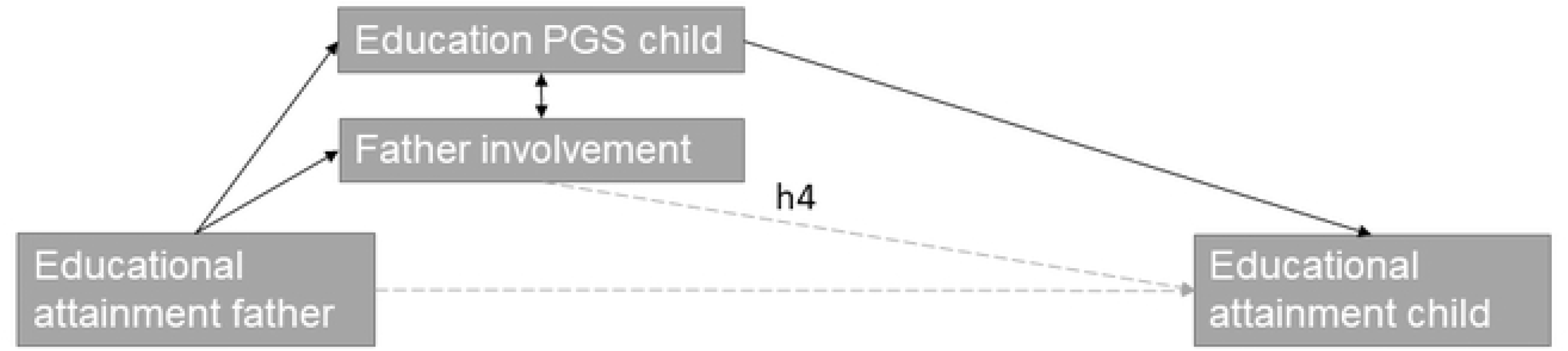
Graphical presentation of hypothesis 4: genetic confounding.

### Implications of correlations between genetic factors and father involvement for understanding the mechanisms underlying the intergenerational transmission of educational attainment: Social confounding

Alternatively, it might be the case that some of the effects we find for children’s education PGS on children’s educational attainment are confounded by the environment, in our case father involvement (see Figure 3 for a graphical presentation of this hypothesis). The rationale for the existence of social confounding is that the Education PGS is based on the GWAS of a large number of SNPs that are associated with educational attainment. Yet, these associations do not imply direct causation, and the pathways from genetic variants to education are diverse. GWAS studies that are used to create PGSs cannot distinguish between “direct” genetic effects-associations between genes and education through intelligence and motivation- and indirect genetic effects -associations between genes and education due to the family environment and parenting practices. As such, part of the effect of our genetic factors might be driven by our behavioural factors; children’s education PGS is associated with children’s educational attainment, because children’s education PGS is associated with the parenting of the child’s father, which is driven by his Education PGS, and it is father involvement that is shaping children’s educational attainment. Previous research found that the effect of mother’s genes on children’s education is partly explained by mothers’ cognitively stimulating parenting (8) and that the effect of children’s own genes on their education is reduced once family SES and household chaos are taken into account (10). Studies also showed that the genes that parents do not transmit to their children impact their children’s educational attainment, which is likely due to the environment parents provide to their children(49–51). This hints towards the idea that also the genes that parents do transmit to their children partly influence their children through the home environment. Based on the above, we expect that *father involvement partly explains the genetic mechanism underlying the intergenerational transmission of education (H5)*.

**Fig 3.**
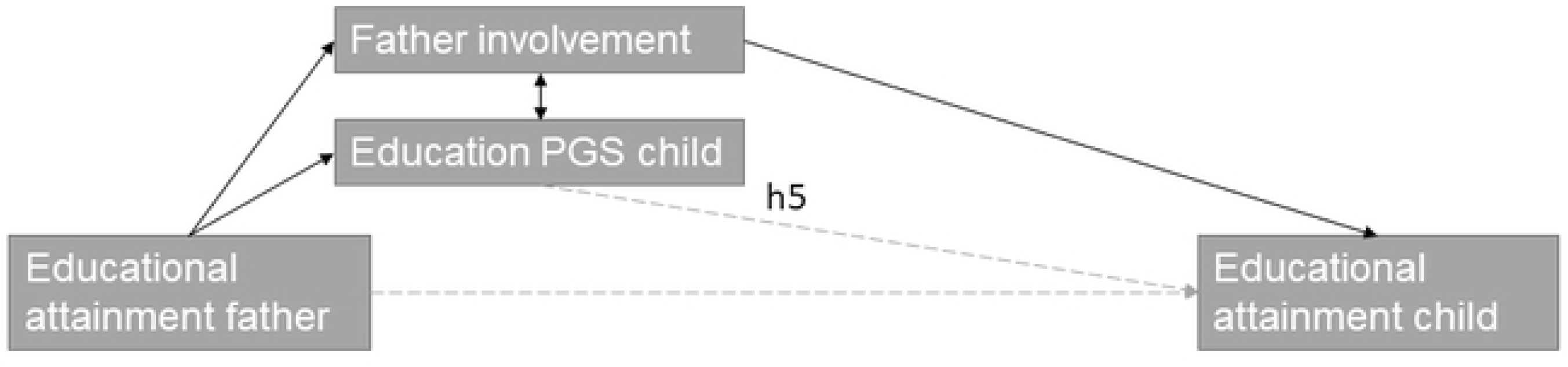
Graphical presentation of hypothesis 5: social confounding.

### Zooming in: Understanding which type of gene-environment correlation is at play

Previous research has not always been able to differentiate passive rGE from active and evocative correlations, which is unfortunate if one wants to understand why this correlation exists. In our study, we can differentiate active/evocative from passive rGE, by using a subsample of our data that includes sibling data. The presence of within-family correlations between children’s education PGS and father involvement (i.e. differences in the education PGS of siblings correlate with differences in father involvement towards these siblings) would imply active/evocative rGE. However, if within families we find a substantially smaller correlation between father involvement and children’s genes as compared to between-families, this would indicate that the correlation is mainly driven by characteristics of the family, which is indicative of passive rGE. Although no previous studies examined passive versus active rGE regarding father involvement, research found support for child evoked rGE, showing that children’s genotype evokes parental warmth, sensitivity, harsh discipline, negative effect (48), but also passive rGE has been found for example regarding warmth, sensitive and stimulating parenting (8,9,52). We expect to find *passive rGE for both indicators of father involvement (father’s school-specific involvement and leisure involvement) (H6).* In addition, given that father involvement might be explicitly activated or evoked based on children’s genetic disposition towards school, we expect that *child-evoked/active rGE will only be found for father’s school-specific involvement (H7)*.

### Current study

This study aims to gain a better understanding of the mechanisms underlying the intergenerational transmission of educational attainment by scrutinizing the interrelatedness of genetic influences and father involvement. We will answer 5 related research questions: 1) To what extent do genes and father involvement independently explain the intergenerational transmission of education, 2) to what extent do genes and behaviour correlate in their relation to educational attainment, 3) to what extent do genes explain part of the behaviour mechanism (‘genetic confounding’), 4) to what extent does behaviour explain part of the genes mechanism (‘social confounding’) and 5) to what extent can we distinguish between passive and active/evocative rGE? We will examine these research questions using data from the National Longitudinal Study of Adolescent to Adult Health (Add Health) of 5,021 genotyped individuals. These data contain information on father involvement, as well as the same information for mothers. Controlling for maternal involvement is important, as this allows us to isolate the independent effects of paternal involvement. Due to amongst others behavioural contagion, in which parents copy each other’s behaviour, mothers and fathers within the same household are more likely to show similar behaviours (e.g. Barnett, Deng, Mills-Koonce, Willoughby, & Cox, 2008; Guay, Ratelle, Duchesne, & Dubois, 2018) also concerning parental involvement (55). Because our sample of respondents has been genotyped, we can include the child’s education PGS, which captures the additive effect of common genetic influences on education. In addition, using sibling data, available for a subsample within our data, we can tap into the *causes* of the hypothesized correlation between father involvement and education PGS. These sibling data allow us to unravel whether observed correlations between father involvement and education PGS are attributable to passive versus active/evocative gene-environment correlations, which have not been assessed yet for father involvement. Respondents in our sample are followed from age 12-19 up to age 32-42.

## Methods

### Data

The Add Health data is a longitudinal study of adolescents from the United States who were in grades 7 to 12 in wave 1 in 1994/95 (56). The data were collected using a school-based stratified sample, in which 80 high schools in the United States were selected, and from these 80 high schools, 20,745 adolescents in grades 7 to 12 participated (56). In this first wave, respondents were between 12 and 19 years old, with the majority between 14 and 18 years old. In the second wave, in 1996, respondents were between 13 and 21 years old. In the third wave, in 2001/02, respondents were between 18 and 26. At wave 4 in 2008/09, respondents were between 24 and 32. Finally, during wave 5 in 2016/18, respondents were between 32 and 42 years old.

From the 20.745 respondents, we selected respondents who were followed up to wave 4 or 5, since these respondents are between the ages 24 and 42 and were thus most likely to have finished their educational training. This restriction reduced our sample size to 17,535. After this, we selected only respondents who reported the educational attainment of their biological father and biological mother, which narrowed down our sample size to 14,712. After this, we only included respondents with an education PGS, which reduced the sample to 5,021 individuals. Finally, we removed individuals with missing information on parental involvement, resulting in a final sample size of 4,579 respondents.

*Genotyping* In total 96% of the participants in wave 5 gave saliva, and 80% of them consented with long-term archiving of their data and were eligible for genome-wide genotyping (57). This resulted in approximately 12,200 genotyped respondents. After quality control, data were available for 9,974 individuals (please see Highland et al (2018) for details on quality control). Non-European descent individuals were removed from the sample, as PGSs based on a GWAS with European-descent individuals is much less suitable for non-European descent individuals, leaving 5,787 individuals. After additional quality control, 5,690 individuals remained with valid genetic information.

Individuals were genotyped using the Illumina Omni1-Quad BeadChip (80% of the sample) and the Illumina Omni2.5-Quad BeadChip (20% of the sample). Before quality control, 609,130 SNPs were available. After quality control (call rate <0.98, HW p-value <10^-4^, MAF<0.01) 346,754 SNPs remained and were used for imputation. Imputation was done using the Haplotype Reference Consortium (HRC) v1.1 European reference panel.

Sibling data: The Add Health contains in total 3,139 sibling pairs, consisting of full-siblings, half-siblings, and MZ and DZ twin pairs and unrelated siblings. Reducing this sample to only sibling pairs with education PGS available reduces the sample size to 619 sibling pairs, of which 429 full siblings and DZ twin pairs, which is further reduced to 380 sibling pairs once we only include those who have information of father involvement available.

### Ethical considerations

The Medical Ethics Review Board of the Erasmus Medical Center Rotterdam considered whether or not this research falls within the scope of the Medical Research Involving Human Subjects Act (WMO). It was concluded that the research is not a clinical research with test subjects as meant in the Medical Research Involving Human Subjects Act (WMO). Therefore, the Medical Ethics Review Board of the Erasmus Medical Center Rotterdam had no task in reviewing the protocol. Therefore it was concluded that we were allowed to conduct the research. Add Health participants provided written informed consent for participation in all aspects of Add Health.

### Measures

#### Years of education of the child

In waves 4 and 5 respondents were asked about the highest level of education that they completed. Answer categories ranged from ‘8^th^ grade or less’, to ‘completing a post-baccalaureate professional degree’. We recoded this variable to years of education. Response options and the years of education that correspond to them (in parentheses) were: 8th grade or less (8), some high school (10), high school graduate (12), some vocational/technical training (13), completed vocational/ technical training (14), some college (14), completed college (16), some graduate school (17), completed a master’s degree (18), some graduate training beyond a master’s degree (19), completed a doctoral degree (20), some post-baccalaureate professional education (18), and completed post-baccalaureate professional education (19). This is in line with previous studies that coded years of education in AddHealth (26,59,60). To reduce missing data, for those respondents who did not give information in wave 5, we used the information from wave 4.

#### Years of education of parent

In wave 1, respondents were asked about the educational attainment of their biological parents. Answer categories ranged from ‘8^th^ grade or less’, to ‘professional training beyond a four-year college or university’. We recoded this variable to years of education. Response options and the years of education that correspond to them (in parentheses) were: never went to school (0), 8th grade or less (8), more than eighth grade, but did not graduate from high school (10), went to a business, trade, or vocational school instead of high school (10), high school graduate (12), completed a GED (12), went to a business, trade, or vocational school after high school (14), went to college, but did not graduate (14), graduated from a college or university (16), professional training beyond a four-year college or university (18). For a subset of the respondents, parents provided information on their educational attainment themselves. Our robustness checks show that the correspondence between child reports and parent reports is high: correlation of 0.87 for mothers and 0.79 for fathers, see Supplementary Material part 3).

#### Father’s school-specific involvement

In wave 1, respondents were asked if, in the past four weeks, they talked with their father about schoolwork or grades (yes or no), worked with their father on a project for school (yes or no), and talked with their father about other things they are doing in school (yes or no). These measures were added up, ranging from 0 for children who did none of these school-related activities with their father to 3 for those who did all these activities.

#### Father’s leisure involvement

In wave 1, respondents were asked if in the past four weeks they went shopping with their father (yes or no), played sports with their father (yes or no), talked with their father about someone they’re dating or a party they went to (yes or no), went to a movie, play, museum, concert or sports event with their father (yes or no) or talked about a personal problem with their father (yes or no). These activities were added up, ranging from 0 if the respondent did none of these activities with their father, to 5, if they did all these activities with their father.

#### Polygenic score (PGS) for years of education

We include PGS for years of education based on the GWAS conducted among 1.1 million individuals (26). The PGS was constructed by the Social Science Genetic Association Consortium (SSGAS) (57) and provided by Add Health. This PGS was created using LDpred, which uses all SNPs and weights them according to their conditional effect, given all other SNPs.

### Controls

*Father residence* We distinguish between whether the child lived with the biological father at wave 1 (resident father=1) or not (non-resident father=0).

*Principal components* To control for population stratification, which is the case if certain SNPs are more common in certain ancestry groups than others, we control for the first 10 principal components (PCs). These PCs were created by the SSGAS (57).

In our models, we control for the *age the respondent had at the first interview*, which ranged between 12 and 21, with the majority of the respondents between 14 and 18. We control for the *sex of the respondent*, and whether or not the respondent was *enrolled in school* at the last wave of data collection.

To examine the unique contribution of father involvement, we control for certain characteristics of the mother, namely *years of education of the mother* and maternal involvement, namely *mother’s school-specific involvement*, *mother’s leisure involvement*, and whether or not the *child lived with the biological mother at wave 1*.

### Analyses

#### Path model to test for mediation (hypothesis 1 and 2) and confounding (hypothesis 4 and 5)

To estimate the extent to which fathers’ involvement and children’s PGS for years of education mediated the relationship between fathers’ years of education and children’s years of education (hypothesis 1 and 2), a path model was estimated using the Lavaan package in R (61). Since the respondents in our sample are not independent but nested within households, we ran a multilevel path model of individuals nested within households (62). A bootstrap sampling procedure was used to estimate the significance of the indirect effects (63).

In our path model, we simultaneously tested the direct effect of the father’s years of education on the years of education of his child, as well as the mediated effect via father’s involvement and the child’s PGS for years of education. These mediated effects are estimated by multiplying the coefficient of (a) the independent variable on the mediators and (b) the mediators on the outcome (64).

Because we wanted to assess potential confounding of genetic effects by father involvement and vice versa (hypothesis 4 and 5), three nested path models were fitted, first a multiple mediation model in which only the two aspects of father involvement are assessed simultaneously, second a mediation model in which the role of children’s PGS for years of education is examined, and third a multiple mediation model that includes both father involvement and children’s education PGS. To quantify the extent to which the effect of father involvement in explaining the intergenerational transmission of education is confounded by the education PGS, we compare the coefficients of father involvement between the first model (in which only father involvement is included as a mediator) and the third model (in which also the education PGS is included) (64). The other way around, to quantify the extent to which the effect of the education PGS is partly socially confounded, and can be explained by father involvement, we compare the coefficient of the education PGS between the second model (in which only the education PGS is included as a confounder), and the third model (in which both the education PGS and father involvement are included).

#### rGE (hypothesis 3, 6, and 7)

To examine to what extent the education PGS and father involvement correlate, we estimated the correlation between children’s years of education predicted from the PGS model and children’s years of education predicted from the father involvement model. To this end, we first regressed children’s years of education on both parents’ years of education, the first 10 PCs, and all other control variables using a multilevel model that takes into account the nested structure of the data. The residuals from this model were used and regressed on the education PGS (model 1), father’s school-specific involvement (model 2), and father’s leisure involvement (model 3). Finally, we assessed the correlation between the predicted values from model 1 with model 2 and model 3.

To examine active/evocative and passive rGE, we estimate rGE between and within families. Between families, we cannot simply examine the correlation between father involvement and the education PGS, as we not only have to control for spurious associations based on ancestral differences but also because we have to take the nested structure of the data into account. Therefore, we estimated multilevel regression models in which we explain the two measures of father involvement by the education PGS while controlling for the first 10 PCs (hypothesis 3).

To assess the rGE within families, we do not have to control for the first 10 PCs, as siblings share their ancestry, and we do not have to take into account the nested structure of the data as there is only one sibling pair per family. Therefore, we estimated linear regression models in which we explain the difference in father involvement between siblings by the difference in the education PGS between siblings. To distinguish between active and passive rGE (hypotheses 6 and 7), we will compare estimates of rGE within and between families.

## Results

### Univariate descriptives

Sample descriptives can be found in Table 1. The respondents in our sample finished 14.63 years of education on average, which equals some years of education after finishing high school (12 years equals finishing high school and 16 years equals finishing a bachelor’s degree). Their parents on average were in school a bit shorter (on average 13.5 years). Fathers are on average less involved with their children’s school activities and leisure than mothers were. Children’s age at the first interview was on average 16 and they were almost 36 on average at the final wave when they were asked about their final education. 71% lived with their father at the first wave, and 29% did not, while a little under 10% did not live with their mother at the time of the first wave of data collection.

**Table 1.** Univariate descriptives of the sample

|  | Mean | SD | Min | Max |
| --- | --- | --- | --- | --- |
| Years of education child | 14.62 | 2.32 | 8 | 20 |
| Years of education father | 13.52 | 2.54 | 0 | 18 |
| Years of education mother | 13.48 | 2.35 | 0 | 18 |
| PGS education child | 0.55 | 0.15 | 0 | 1.05 |
| Father's school-specific involvement | 1.16 | 1.01 | 0 | 3 |
| Father's leisure involvement | 1.44 | 1.29 | 0 | 5 |
| Mother's school-specific involvement | 1.32 | 1 | 0 | 3 |
| Mother's leisure involvement | 2.08 | 1.21 | 0 | 5 |
| Age first interview | 16.01 | 1.7 | 12 | 21.33 |
| Age asked education | 35.83 | 4.13 | 25 | 43 |
|  | Yes n | % | No n | % |
| Enrolled in educ last wave | 355 | 7.75 | 4223 | 92.25 |
| Live with father at the first wave | 3427 | 74.84 | 1152 | 25.16 |
| Live with mother at the first wave | 4181 | 91.31 | 398 | 8.69 |

### Bivariate descriptives

Figure 4 shows the correlation between the variables in our analyses. Children’s years of education is positively correlated (around 0.4) with the education of both parents and children’s Education PGS. Furthermore, both father’s and child’s years of education are positively correlated with father’s school-specific involvement and father’s leisure involvement. The child’s education PGS is positively correlated with father’s school-specific involvement and father’s leisure involvement. There is also a positive correlation between the two different aspects of father involvement (father’s school-specific involvement and father’s leisure involvement).

**Fig 4.**
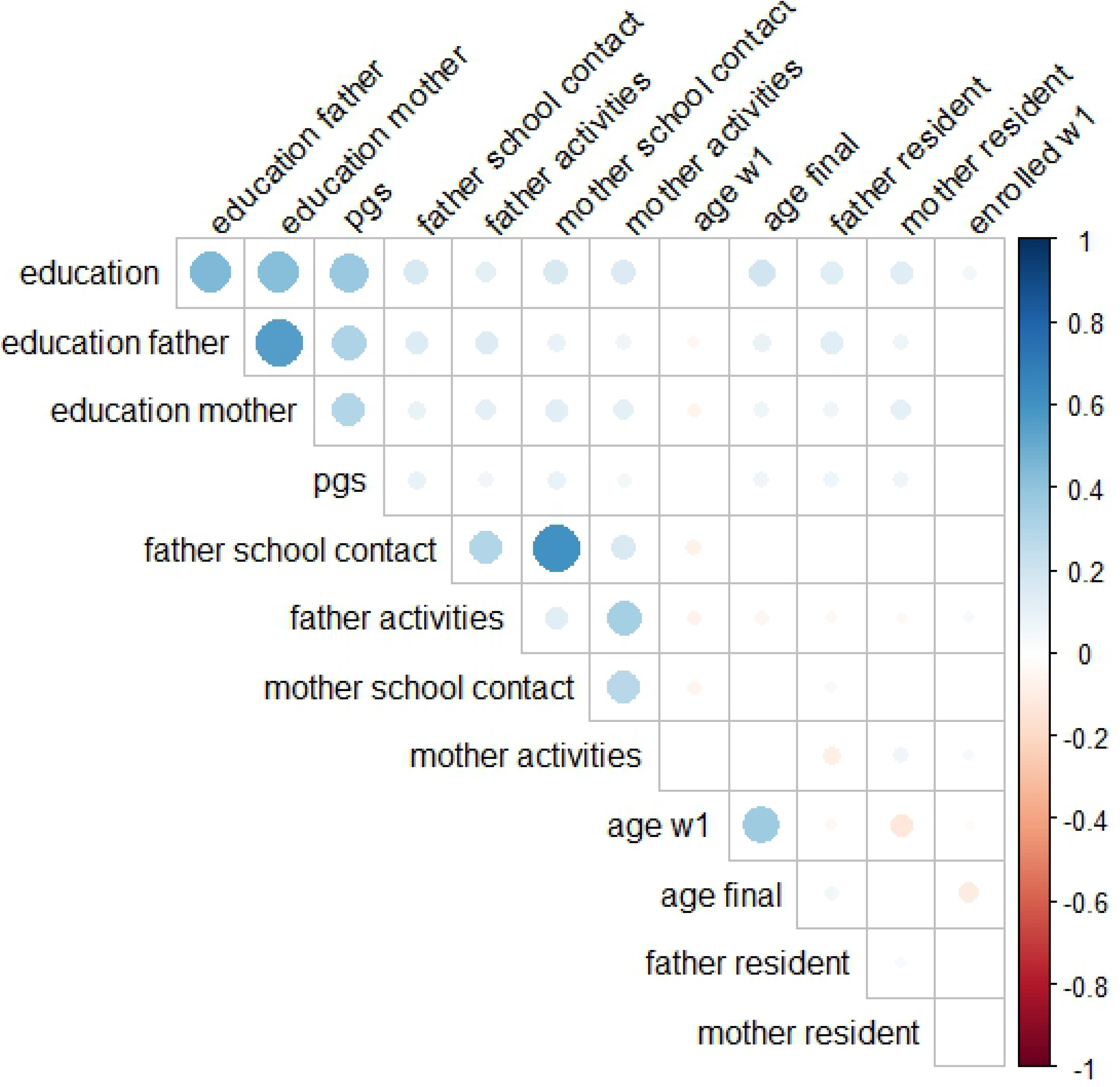
Correlation table graphically displayed. Non-significant correlations are not displayed. The size and color represent the strength and direction of the correlation. Education=Years of education child; Education father=Years of education father; Education mother=Years of education mother; pgs=Child’s Polygenic score for education; father school contact=father’s school-specific involvement; father activities=father’s leisure involvement; mother school contact=Mother’s school-specific involvement; mother activities=mother’s leisure involvement; age w1= age at wave 1; age final=age at the final wave; father resident= lived with father at wave 1, 0= no 1=yes; mother resident= lived with mother at wave 1, 0= no 1=yes; enrolled w1: was the respondent enrolled in education at wave 1, 0=no, 1=yes.

### Part 1) Regression results: mediation

Our first two hypotheses focused on the extent to which the association between fathers’ years of education and children’s years of education is mediated by father involvement (hypothesis 1) and the child’s education PGS (hypothesis 2). To this end, we estimated path models in which we examined the direct effect of education fathers and the significance of the indirect effects via father involvement and education PGS. The results are displayed in Table S1 and Figure 5. Father’s school-specific involvement significantly mediates 2.3% of the intergenerational transmission (0.007 of the total effect of 0.303), father’s leisure involvement 1.3% (0.004 of 0.303), and the education PGS significantly mediates 21.45% (0.065 of 0.303). These findings support both hypotheses 1 and 2 and show that both genes and father involvement are significant mediators. In addition, our findings show that the education PGS mediates a much larger proportion of the intergenerational transmission of years of education than our two measures of father involvement.

**Fig 5.**
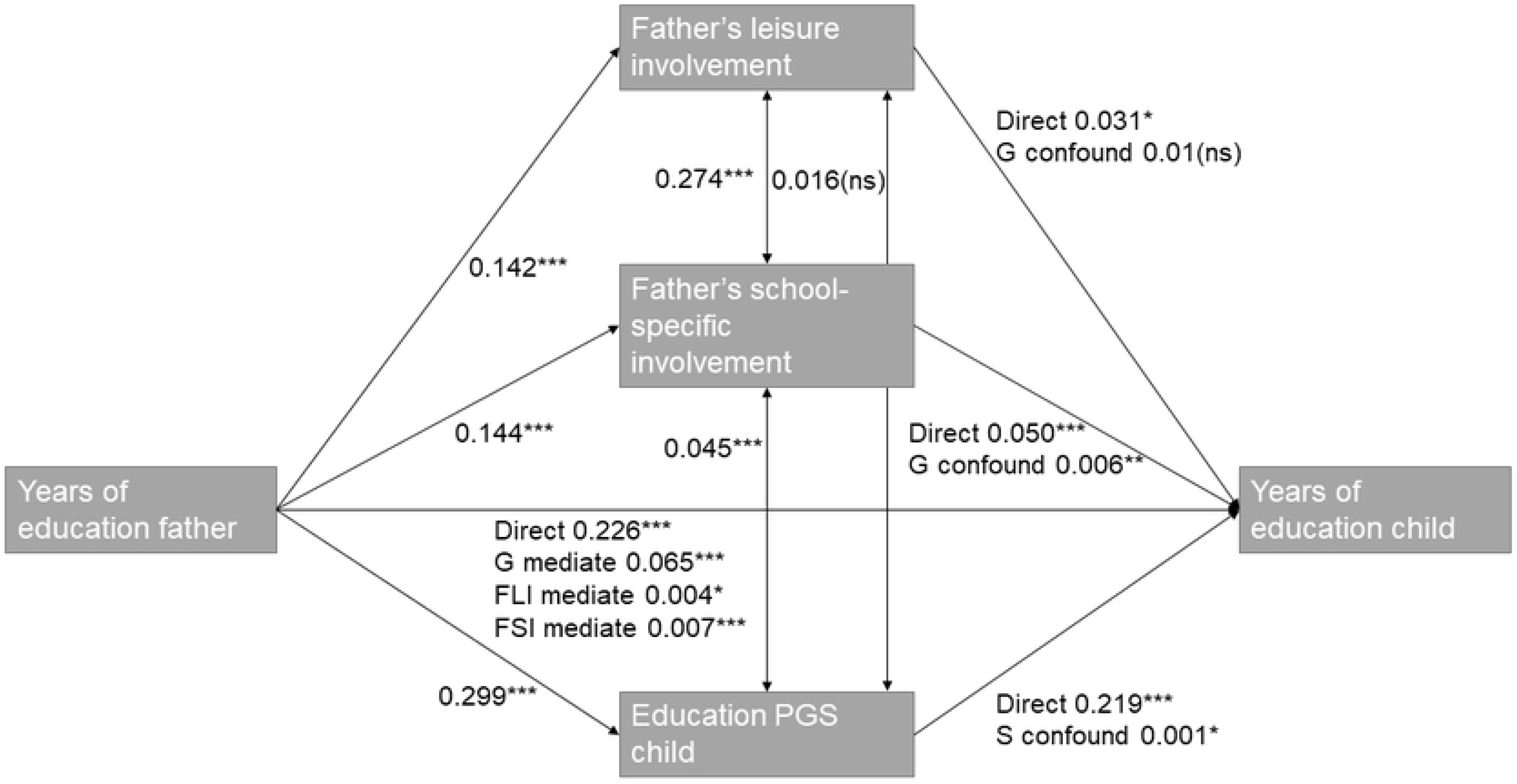
Graphical presentation of the mediation and confounding results. G mediate refers to the part of the effect that is genetically mediated, FLI mediate refers to the mediated effect by father’s leisure involvement, FSI mediate refers to the mediated effect by father’s school-specific involvement, G confound to the genetically confounded part and S confound to the social confounding by father involvement.

### Part 2) rGE

Table 2 displays the correlations between the child’s education PGS and our two dimensions of father involvement. The correlated effects are displayed in the left panel of Table 2. We find significant correlated effects between father’s school-specific involvement and the education PGS of 0.092, and somewhat smaller positive significant correlated effects between father’s leisure involvement and the education PGS of 0.063 (see Table 2 left panel).

**Table 2.**
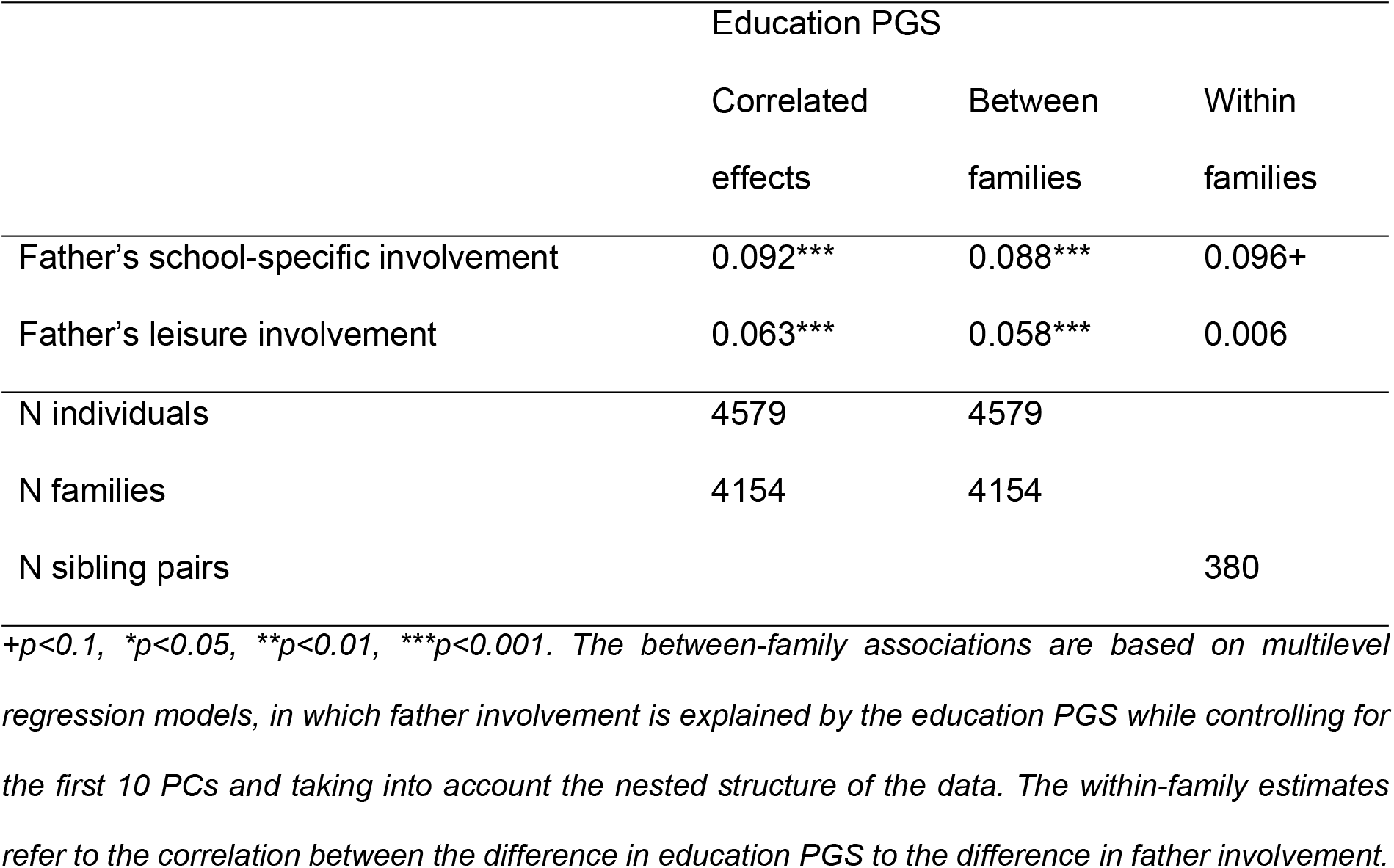
Correlations between education PGS and father involvement: correlated effects, between-family, and within-family correlations

|  | Education PGS |  |  |
| --- | --- | --- | --- |
|  | Correlated effects | Between families | Within families |
| Father's school-specific involvement | 0.092*** | 0.088*** | 0.096+ |
| Father's leisure involvement | 0.063*** | 0.058*** | 0.006 |
| N individuals | 4579 | 4579 |  |
| N families | 4154 | 4154 |  |
| N sibling pairs |  |  | 380 |

+ $p < 0.1$ , \* $p < 0.05$ , \*\* $p < 0.01$ , \*\*\* $p < 0.001$ . The between-family associations are based on multilevel regression models, in which father involvement is explained by the education PGS while controlling for the first 10 PCs and taking into account the nested structure of the data. The within-family estimates refer to the correlation between the difference in education PGS to the difference in father involvement.

### Part 3) Genetic confounding

Thirdly, we were interested in genetic confounding; the extent to which accounting for genetic influences reduces the effect of father involvement. Results are displayed in Figure 5. The effect of the father’s school-specific involvement is significantly reduced from 0.056 to 0.050 when including the child’s education PGS, which implies 10.7% genetic confounding, while the effect of the father’s leisure involvement is not significantly reduced when including children’s education PGS. The former shows that a substantial part of the role the father’s school-specific involvement plays in the intergenerational transmission of years of education is genetically confounded, which supports hypothesis 4. The latter shows that the role the father’s leisure involvement plays in the intergenerational transmission of years of education is not genetically confounded, which contrasts with hypothesis 4.

### Part 4) Social confounding

Furthermore, we were interested in social confounding: the extent to which the role of children’s genes in the intergenerational transmission of years of education is reduced when information on father involvement is included in the model. Results are again displayed in Figure 5. Our results showed that, once father involvement is added to the model, the role of children’s education PGS as the underlying mechanism is only slightly reduced, from 0.220 to 0.219, which implies a mere 0.5% social confounding. Furthermore, this significant social confounding only holds for the father’s school-specific involvement. These findings support hypothesis 5 but are very small in magnitude.

### Part 5) rGE within versus between families

Comparing the between-families and within-families associations between father involvement and children’s education PGS provides insights into the extent to which the rGE is active or passive. For father’s school-specific involvement, we find suggestive evidence of within-family correlation (0.096+, borderline significant), and a significant between-family correlation (0.088***), see Table 2. This borderline significant within-family correlation suggests that the sibling with the highest Education PGS also reports more father’s school-specific involvement. This hints towards child-evoked rGE, as the child with the highest education PGS behaves in a certain way that evokes involvement of the father, or active rGE, as the child with the highest education PGS actively involves his/her father in their schoolwork. For the father’s leisure involvement, we only find between-family correlation (0.058***), implying passive rGE from the side of the child. These findings are in line with hypothesis 6, showing passive rGE, and show suggestive evidence of hypothesis 7 on active rGE for father’s school-specific involvement.

### Robustness checks

We conducted several additional analyses to assess the robustness of our findings. To assess the robustness of our findings of the effect of the education PGS, we tested if this effect was inflated by indirect effects, due to population stratification, or due to assortative mating. We found that the within-family effect of the education PGS was smaller than the between family effect, yet it remained significant, which shows that controlling for indirect effects and population stratification does not fully explain our findings regarding the association between the education PGS and years of education. Our findings indicate that assortative mating could to a small extent result in an overestimation of the effect of the education PGS (see Robustness checks S1). We furthermore assessed whether the association between our measures of father involvement and education depends on the age of the respondents during the first interview. Our robustness checks reveal it does not (see Robustness checks S2). We also assessed whether any multicollinearity issues arise when father involvement and mother involvement were added simultaneously to the model. Our analyses revealed no multicollinearity issues (see Robustness checks S2). Finally, we examined the robustness of our findings by assessing whether our results differ when we use a parent-report or child-report on parental education. Our analyses revealed that our conclusions are similar, irrespective of the reported used (see Robustness checks S3).

## Conclusion and discussion

This study aimed to gain a better understanding of the mechanisms, and their interrelatedness, underlying the intergenerational transmission of educational attainment. To this end, we answered five interrelated questions.

Our first question was to examine whether and to what extent father involvement and children’s education PGS were unique and independent mechanisms underlying this intergenerational transmission of educational attainment. Our results revealed that both dimensions of father involvement as well as children’s education PGS were significant and unique mechanisms. Noteworthy is that our results indicated that children’s education PGS was a much more important underlying mechanism than the two dimensions of father involvement we considered in the current study. Our relatively small effects of the family environment compared to the genetic component is in contrast to findings from Allegrini and colleagues (10), who found that multiple PGSs can explain 15% of the variance in educational attainment, while environmental factors can explain 28%. The higher percentage for environmental factors found in their study is likely attributable to the fact that they used a much broader indication of the family environment, consisting of amongst others the educational attainment of both parents, employment status of parents, chaos at home, and important life events.

Our second question focused on the extent to which children’s education PGS and father involvement correlate as mechanisms underlying the intergenerational transmission of educational attainment. Our results indicated that both mechanisms indeed correlate. These findings add to the previous research that shows a correlation between other aspects of parenting and the education PGS, such as parental warmth and sensitivity (9). This correlation implies that many children are either growing up with double disadvantages – having both a less involved father and a lower education PGS – or double advantages – a highly involved father and a higher education PGS. This highlights the importance for social scientists to take genetic influences into account when examining the importance of parenting in the intergenerational transmission of education, and for genetic scientists to take into account social pathways.

Our third question centred on genetic confounding; to what extent is the role that father involvement plays in the intergenerational transmission of educational attainment explained by children’s education PGS. Our results indicated that genetic confounding plays a substantial role when it comes to the role fathers’ school-specific involvement plays as an underlying mechanism in the intergenerational transmission of educational attainment. This indicates that findings from the field of behavioural sciences have likely overestimated the role that fathers’ school-specific involvement plays in the intergenerational transmission of educational attainment. It furthermore suggests that part of “what we think of as measures of ‘environment’ are better described as external factors that might be partly under genetic control” (65). However, and intriguingly, the role of fathers’ leisure involvement as an underlying mechanism hardly changed when information on children’s genes was added to the model. Furthermore, fathers’ school-specific involvement was not fully mediated by the children’s education PGS, indicating that also school-specific involvement independently explains the intergenerational transmission of education. Our findings regarding the relevance of father involvement in the intergenerational transmission of educational attainment suggests that father involvement should be considered as a potential candidate for an intervention to aid in breaking the intergenerational cycle of (dis)advantage. Several interventions, such as educational sessions on parenting skills, as well as policies, such as parental leave reserved for fathers, have been proven to increase father’s involvement (66, 67).

Our fourth question revolved around behavioural confounding; to what extent is the role that children’s education PGS plays in the intergenerational transmission of educational attainment explained by father involvement. Our results indicate that behavioural confounding plays a negligible role. These findings differ from the findings from previous research that examined social confounding of genetic effects (8, 10). Again, these differences are likely attributable to differences in how broadly the family environment was defined.

Fifth, we assessed active versus passive rGE by looking at correlations within and between families. Within families, we find suggestive evidence that the sibling with the highest education PGS also has more contact about school with their father. This could indicate that the correlation between the education PGS and school contact is child evoked or active: the education PGS and the behaviour associated with it (for example motivation or interest in schoolwork) evokes fathers to discuss school with the child or causes the child to actively involve their father in their schoolwork. However, given that this correlation is only marginally significant, future research with larger sibling samples should replicate these analyses. The correlation between the education PGS and activities between father and child is more passive from the perspective of the child, given that we do not find a within-family correlation. This indicates that children with a high education PGS grow up in families with highly educated fathers/fathers with a high education PGS, who have a more active parenting style and therefore perform more activities with their child.

Some limitations need to be considered when interpreting these findings. First, we believe that our measures of father involvement are relevant and cover important dimensions of involvement of fathers, yet there are other aspects of father involvement that are also relevant that we did not cover. Our measures largely tap into the quantity of involvement (how much do fathers and children discuss school matters and undertake activities together). Other studies have emphasized that pertaining to children’s educational outcomes, the quality of parental involvement is also important (68). It is therefore likely that the current study only yielded an underestimation of the role father involvement plays in the intergenerational transmission of educational attainment.

Similarly, our genetic measure only captures a part of the genetic component of education, as twin studies showed a heritability of 40% (6) while the education PGS explains approximately 10%. This discrepancy between relatively high heritability estimates from twin studies and lower explained variance from PGSs has been called missing heritability and has been found for many traits (69). Possible reasons are amongst others that PGSs only captures common genetic variants and only additive effects, while rare variants and non-additive effects might be relevant as well. Therefore, our estimates of genetic mediation of the intergenerational transmission of education, and genetic confounding of father involvement, are underestimations and only capture part of the genetic effect

When interpreting our findings regarding father involvement, we must consider that we study the role of fathers in the United States in 1994. In this time period involvement of fathers was much lower than in more recent cohorts (70). The greater involvement of fathers in more recent cohorts could imply that fathers play an even bigger role in the educational achievements of their children, and therefore in the intergenerational transmission of education in these cohorts. However, the group of fathers that exhibited strong involvement in their children’s lives was most likely more selective in older cohorts than in recent ones. Consequently, it might also be the case that father involvement played a less substantial role as underlying mechanism in more recent cohorts than in older ones.

Our study is limited by the fact that our sample only includes respondents of European ancestry (i.e. White respondents). The reason is that the GWAS for educational attainment is based only on European ancestry individuals, and PGSs created from such GWASs have lower predictive power among non-European samples (71). Therefore, we cannot generalize our findings to other ancestry groups. For this reason, we also cannot use our findings to explain differences in educational achievement between different ancestry groups.

To summarize, we find that both genes and father involvement are underlying mechanisms in the intergenerational transmission of educational attainment, that these mechanisms are correlated with each other, and that part of the role that fathers’ school-specific involvement plays as underlying mechanism is confounded by children’s education PGS. Our findings underscore the need to control for genetic effects in studies that examine the role of parenting in the intergenerational transmission of inequality, but also the need to control for parental involvement and the family environment in general when considering the role that genes play in this intergenerational transmission. Thus, our study underscores that in order to fully understand the mechanisms that underly intergenerational reproduction of (dis)advantages, scholars need to integrate both insights and data from different disciplines.

## Supporting information

**S1 Robustness checks regarding the education PGS**

**S2 Robustness checks regarding father involvement**

**S3 Education of parent reported by the respondent and by the parent**

**Table S1** Results from the multilevel path model on years of education child

**Table S2** Effect of the Education PGS on years of education in the 1) OLS model with only PCs as controls, 2) OLS model including PCs and additional controls and 3) Fixed Effects model

**Table S3** Correlation between age of the respondent at the first wave, and measures of father involvement

**Table S4** Mean and SD of measures of father involvement for the different ages

**Figure S1** Association between education PGS and years of schooling in the complete sample

**Figure S2** Association between the difference in education PGS and difference in years of education between siblings.

## Acknowledgements

The present study was supported by a grant from the Netherlands Organization for Scientific Research to RK (NWO MaGW VIDI; grant no. 452-17-005) and by a grant from the European Research Council to RK (ERC StG; grant no. 757210). Both authors declare no conflict of interest.

Add Health is directed by Robert A. Hummer and funded by the National Institute on Aging cooperative agreements U01 AG071448 (Hummer) and U01AG071450 (Aiello and Hummer) at the University of North Carolina at Chapel Hill. Waves I-V data are from the Add Health Program Project, grant P01 HD31921 (Harris) from Eunice Kennedy Shriver National Institute of Child Health and Human Development (NICHD), with cooperative funding from 23 other federal agencies and foundations. Add Health was designed by J. Richard Udry, Peter S. Bearman, and Kathleen Mullan Harris at the University of North Carolina at Chapel Hill.

## References

1. Pfeffer FT. Persistent inequality in educational attainment and its institutional context. Eur Sociol Rev. 2008;24(5):543–65.

2. Passaretta G, Skopek J. Roots and Development of Achievement Gaps. A Longitudinal Assessment in Selected European Countries. Dublin; 2018. (ISOTIS Report). Report No.: D1.3.

3. OECD. Education at a Glance 2017 [Internet]. Education at a Glance 2017. 2017. 25– 26 p. Available from: https://doi.org/10.1787/eag-2017-en

4. Keizer R, Van Lissa CJ, Tiemeier H, Lucassen N. The Influence of Fathers and Mothers Equally Sharing Childcare Responsibilities on Children’s Cognitive Development from Early Childhood to School Age: An Overlooked Mechanism in the Intergenerational Transmission of (Dis)Advantages? Eur Sociol Rev. 2020;36(1):1–15.

5. Lareau A. Invisible inequality: Social class and childrearing in black families and white families. Am Sociol Rev. 2002;67(5):747–76.

6. Branigan AR, McCallum KJ, Freese J. Variation in the Heritability of Educational Attainment: An International Meta-Analysis. Soc Forces [Internet]. 2013 Jun 25 [cited 2014 Nov 11];92(1):109–40. Available from: http://sf.oxfordjournals.org/cgi/doi/10.1093/sf/sot076

7. Krapohl E, Hannigan LJ, Pingault J-B, Patel H, Kadeva N, Curtis C, et al. Widespread covariation of early environmental exposures and trait-associated polygenic variation. Proc Natl Acad Sci [Internet]. 2017;201707178. Available from: http://www.pnas.org/lookup/doi/10.1073/pnas.1707178114

8. Wertz J, Moffitt TE, Agnew-Blais J, Arseneault L, Belsky DW, Corcoran DL, et al. Using DNA from mothers and children to study parental investment in children’s educational attainment. Child Dev. 2020;91(5):1745–61.

9. Wertz J, Belsky J, Moffitt TE, Belsky DW, Harrington H, Avinun R, et al. Genetics of nurture: A test of the hypothesis that parents’ genetics predict their observed caregiving. Dev Psychol [Internet]. 2019; Available from: http://doi.apa.org/getdoi.cfm?doi=10.1037/dev0000709

10. Allegrini AG, Karhunen V, Coleman JRI, Selzam S, Rimfeld K, Stumm S Von, et al. Multivariable G-E interplay in the prediction of educational achievement. PLoS Genet [Internet]. 2020;1–20. Available from: http://dx.doi.org/10.1371/journal.pgen.1009153

11. Bates TC, Maher BS, Medland SE, McAloney K, Wright MJ, Hansell NK, et al. The Nature of Nurture: Using a Virtual-Parent Design to Test Parenting Effects on Children’s Educational Attainment in Genotyped Families. Twin Res Hum Genet. 2018;21(2):73–83.

12. Hook JL. Care in context: Men’s unpaid work in countries, 1965-2003. Am Sociol Rev. 2006;71(august):639–60.

13. Yeung WJ, Sandberg JF, Davis-kean PE, Hofferth SL. Children’s Time With Fathers in Intact Families. J marriage Fam. 2010;63(1):136–54.

14. Cabrera NJ, Tamis-lemonda CS, Bradley RH, Hofferth S, Lamb ME. Fatherhood in the Twenty-First Century. 2000;71(1):127–37.

15. Fagan J, Lamb ME, Cabrera NJ. Should Researchers Conceptualize Differently the Dimensions of Parenting for Fathers and Mothers ? J Fam Theory Rev. 2014;6(December):390–405.

16. Kim SW, Hill NE. Including fathers in the picture: A meta-analysis of parental involvement and students’ academic achievement. J Educ Psychol. 2015;107(4):919–34.

17. Altintas E, Sullivan O. Trends in fathers’ contribution to housework and childcare under different welfare policy regimes. Soc Polit. 2017;24(1):81–108.

18. Guryan J, Hurst E, Kearney M. Parental Education and Parental Time with Children. J Econ Perspect [Internet]. 2008;22(3):23–46. Available from: http://pubs.aeaweb.org/doi/10.1257/jep.22.3.23

19. Aughinbaugh A, Robles O, Sun H. Marriage and divorce: patterns by gender, race, and educational attainment. Mon Labor Rev. 2013;

20. Kitterød RH, Lyngstad J. Characteristics of parents with shared residence and father sole custody. Evidence from Norway 2012. 2014.

21. Gordon MS. Self-perception and relationship quality as mediators of father’s school-specific involvement and adolescent’s academic achievement. Child Youth Serv Rev [Internet]. 2017;77(April):94–100. Available from: http://dx.doi.org/10.1016/j.childyouth.2017.04.001

22. Whitney SD, Prewett S, Wang Z, Chen H. Fathers’ Importance in Adolescents’ Academic Achievement. Int J Child, Youth Fam Stud. 2018;8(3/4):101.

23. Nielsen F. Achievement and ascription in educational attainment: Genetic and environmental influences on adolescent schooling. Soc Forces. 2006;85(1):193–216.

24. Johnson W, Deary IJ, Iacono WG. Genetic and environmental transactions underlying educational attainment. Intelligence [Internet]. 2009;37(5):466–78. Available from: http://dx.doi.org/10.1016/j.intell.2009.05.006

25. Schulz W, Schunck R, Diewald M, Johnson W. Pathways of Intergenerational Transmission of Advantages during Adolescence: Social Background, Cognitive Ability, and Educational Attainment. J Youth Adolesc [Internet]. 2017;46(10):2194–214. Available from: http://dx.doi.org/10.1007/s10964-017-0718-0

26. Lee JJ, Wedow R, Okbay A. Gene discovery and polygenic prediction from a genome-wide association study of educational attainment in 1.1 million individuals. Nat Genet. 2018;50:1112–21.

27. Maier RM, Visscher PM, Robinson MR, Wray NR. Embracing polygenicity: a review of methods and tools for psychiatric genetics research. Psychol Med [Internet]. 2017;(May):1–19. Available from: https://www.cambridge.org/core/product/identifier/S0033291717002318/type/journal_article

28. Varghese C, Wachen J. The Determinants of Father Involvement and Connections to Children’s Literacy and Language Outcomes: Review of the Literature. Marriage Fam Rev [Internet]. 2016;52(4):331–59. Available from: http://dx.doi.org/10.1080/01494929.2015.1099587

29. Morales-castillo M. Family Contributions to School Performance of Adolescents : The Role of Fathers ’ Perceived Involvement. J Fam Issues. 2021;(45).

30. Wilder S. Effects of parental involvement on academic achievement: A meta-synthesis. Educ Rev. 2014;66(3):377–97.

31. Pomerantz EM, Moorman EA, Litwack SD. The How, Whom, and Why of Parents ’ Involvement in Children ’ s Academic Lives : More Is Not Always Better. Rev Educ Res. 2007;77(3):373–410.

32. Hill NE, Castellino DR, Lansford JE, Nowlin P, Dodge KA, Pettit GS. Parent Academic Involvement as Related to School Behavior, Achievement, and Aspirations: Demographic Variations Across Adolescence. Child Dev. 2009;75(5):1491–509.

33. Cano T, Perales F, Baxter J. A Matter of Time: Father Involvement and Child Cognitive Outcomes. J Marriage Fam. 2018;81(February):164–84.

34. Huerta MC, Adema W, Baxter J, Han W-J, Lausten M, Lee R, et al. Fathers’ Leave, Fathers’ Involvement, and Child Development: Are They Related? Evidence from Four OECD Countries [Internet]. Paris; 2013. (OECD Social, Employment and Migration Working papers; vol. 88). Report No.: 140. Available from: http://thefamilywatch.org/doc/doc-0077-es.pdf

35. Eccles JS, Harold RD. Family involvement in children’s and adolescents’ schooling. In: Booth A, Dunn JF, editors. Family-school links: How do they affect educational outcomes. 1996. p. 3–34.

36. Hoover-Dempsey K V., Sandler HM. Why do parents become involved in their children’s education? Rev Educ Res. 1997;67(1):3–42.

37. Byford M, Kuh D, Richards M. Parenting practices and intergenerational associations in cognitive ability. Int J Epidemiol. 2012;41(1):263–72.

38. Miller DP, Thomas MMC, Waller MR, Nepomnyaschy L, Emory AD. Father Involvement and Socioeconomic Disparities in Child Academic Outcomes. J Marriage Fam. 2020;82(April):515–33.

39. Matsuoka R, Nakamuro M, Inui T. Emerging inequality in effort: A longitudinal investigation of parental involvement and early elementary school-aged children’s learning time in Japan. Soc Sci Res [Internet]. 2015;54:159–76. Available from: http://dx.doi.org/10.1016/j.ssresearch.2015.06.009

40. Krapohl E, Rimfeld K, Shakeshaft NG, Trzaskowski M, McMillan A, Pingault J-B, et al. The high heritability of educational achievement reflects many genetically influenced traits, not just intelligence. Proc Natl Acad Sci [Internet]. 2014;111(42):15273–8. Available from: http://www.pnas.org/lookup/doi/10.1073/pnas.1408777111

41. Smith-Woolley E, Selzam S, Plomin R. Polygenic Score for Educational Attainment Captures DNA Variants Shared Between Personality Traits and Educational Achievement. J Pers Soc Psychol. 2019;

42. Liu H. Social and Genetic Pathways in Multigenerational Transmission of Educational Attainment. Am Sociol Rev [Internet]. 2018;83(2):278–304. Available from: https://doi.org/10.1177/0003122418759651

43. Ayorech Z, Krapohl E, Plomin R, von Stumm S. Genetic Influence on Intergenerational Educational Attainment. Psychol Sci. 2017;28(9):1302–10.

44. Krapohl E, Plomin R. Genetic link between family socioeconomic status and children’s educational achievement estimated from genome-wide SNPs. Mol Psychiatry [Internet]. 2016;21(3):437–43. Available from: http://dx.doi.org/10.1038/mp.2015.2

45. Plomin R, DeFries JC, Loehlin JC. Genotype-environment interaction and correlation in the analysis of human behavior. Psychol Bull. 1977;84(2):309–22.

46. Ayoub M, Briley DA, Grotzinger A, Patterson MW, Engelhardt LE, Tackett JL, et al. Genetic and Environmental Associations Between Child Personality and Parenting. Soc Psychol Personal Sci. 2019;10(6):711–21.

47. Baizán P, Domínguez M, González MJ. Couple Bargaining or Socio-Economic Status?: Why some parents spend more time with their children than others. Eur Soc [Internet]. 2014;16(1):3–27. Available from: http://dx.doi.org/10.1080/14616696.2013.859717

48. Avinun R, Knafo A. Parenting as a Reaction Evoked by Children’s Genotype: A Meta-Analysis of Children-as-Twins Studies. Personal Soc Psychol Rev. 2014;18(1):87–102.

49. de Zeeuw EL, Hottenga JJ, Ouwens KG, Dolan C V., Ehli EA, Davies GE, et al. Intergenerational Transmission of Education and ADHD: Effects of Parental Genotypes. Behav Genet [Internet]. 2020;50(4):221–32. Available from: https://doi.org/10.1007/s10519-020-09992-w

50. Kong A, Thorleifsson G, Frigge ML, Vilhjalmsson BJ, Young AI, Thorgeirsson TE, et al. The nature of nurture: Effects of parental genotypes. Science (80-). 2018;359(6374):424–8.

51. Hwang L, Tubbs JD, Luong J, Lundberg M, Moen H, Wang G, et al. Estimating indirect parental genetic effects on offspring phenotypes using virtual parental genotypes derived from sibling and half sibling pairs. PLoS Genet [Internet]. 2020;1–29. Available from: http://dx.doi.org/10.1371/journal.pgen.1009154

52. Klahr AM, Burt SA. Elucidating the Etiology of Individual Differences in Parenting: A Meta-Analysis of Behavioral Genetic Research. Psychol Bull. 2014;140(2):544–86.

53. Guay F, Ratelle CF, Duchesne S, Dubois P. Mothers’ and fathers’ autonomy-supportive and controlling behaviors: An analysis of interparental contributions. Parenting. 2018;18(1):45–65.

54. Barnett MA, Deng M, Mills-Koonce WR, Willoughby M, Cox M. Interdependence of Parenting of Mothers and Fathers of Infants. J Fam Psychol. 2008;22(4):561–73.

55. Pleck JH, Hofferth SL. Mother Involvement as an Influence on Father Involvement with Early Adolescents. Fathering. 2008;6(3):1–7.

56. Harris KM, Halpern CT, Whitsel EA, Hussey JM, Killeya-jones LA, Tabor J, et al. Cohort Profile: The National Longitudinal Study of Adolescent to Adult Health (Add Health). Int J Epidemiol. 2019;1–12.

57. Okbay A, Turley P, Benjamin D, Visscher PM, Braudt D, Mullan Harris K. SSGAC Polygenic Scores (PGSs) in the National Longitudinal Study of Adolescent to Adult Health (Add Health). 2018.

58. Highland HM, Avery CL, Duan Q, Li Y, Harris KM. Quality control analysis of Add Health GWAS data. Chapel Hill; 2018.

59. Liu H, Motz RT, Tanksley PT, Barnes JC, Harris KM. Adolescent Criminal Justice Involvement, Educational Attainment, and Genetic Inheritance: Testing an Integrative Model Using the Add Health Data [Internet]. Vol. 7, Journal of Developmental and Life-Course Criminology. Springer International Publishing; 2021. 195–228 p. Available from: https://doi.org/10.1007/s40865-021-00166-8

60. Domingue BW, Belsky DW, Conley D, Mullan Harris K, Boardman JD. Polygenic Influence on Educational Attainment: New evidence from The National Longitudinal Study of Adolescent to Adult Health. AERA Open. 2015;1(3):1–13.

61. Rosseel Y. lavaan : An R Package for Structural Equation. J Stat Softw [Internet]. 2012;48(2):1–36. Available from: http://www.jstatsoft.org/v48/i02

62. Snijders TAB, Bosker RJ. Multilevel Analysis: An introduction to basic and advanced multilevel modeling. 2nd ed. London: Sage publications; 2012.

63. MacKinnon DP, Lockwood CM, Williams J. Confidence limits for the indirect effect: Distribution of the product and resampling methods. Multivariate Behav Res. 2004;39(1):99–128.

64. Mackinnon DP, Krull JL, Lockwood CM. Equivalence of the Mediation, Confounding and Suppression Effect. Prev Sci. 2000;1(4).

65. Vinkhuyzen AAE, van der Sluis S, de Geus EJC, Boomsma DI, Posthuma D. Genetic influences on ‘environmental’ factors. Genes, Brain Behav [Internet]. 2010;9(3):276–87. Available from: http://doi.wiley.com/10.1111/j.1601-183X.2009.00554.x

66. Kalembo FW, Kendall GE. A systematic review of interventions that have the potential to foster engaged fathering to enhance children’s health and development. Child Fam Soc Work. 2021;(November):1–22.

67. Huerta MC, Adema W, Baxter J, Han WJ, Lausten M, Lee R, et al. Fathers’ Leave and Fathers’ Involvement: Evidence from Four OECD Countries. Eur J Soc Secur. 2014;16(4):308–46.

68. Hoskins D. Consequences of parenting on adolescent outcomes. Societies. 2014;4(3):506–31.

69. Manolio TA, Collins FS, Cox NJ, Goldstein DB, Hindorff LA, Hunter DJ, et al. Finding the missing heritability of complex diseases. Nature [Internet]. 2009 Oct 8 [cited 2014 Jul 9];461(7265):747–53. Available from: http://www.pubmedcentral.nih.gov/articlerender.fcgi?artid=2831613&tool=pmcentrez&rendertype=abstract

70. Dotti Sani GM, Treas J. Educational gradients in parents’ child-care time across countries, 1965–2012. J Marriage Fam. 2016;78(4):1083–96.

71. Ware EB, Schmitz LL, Faul J, Gard A, Mitchell C, Smith JA, et al. Heterogeneity in polygenic scores for common human traits. bioRxiv. 2017;(5):1–13.

